# New North American isolates of *Venturia inaequalis* can overcome apple scab resistance of *Malus floribunda* 821

**DOI:** 10.1101/794636

**Authors:** David Papp, Jugpreet Singh, David Gadoury, Awais Khan

## Abstract

Apple scab, caused by *Venturia inaequalis* (Cke.) Wint., is a destructive fungal disease of major apple cultivars worldwide, most of which are moderately to highly susceptible. Thus, development of scab resistant cultivars is one of the highest priorities of apple breeding programs. The principal source of resistance for breeding programs has been the scab resistance gene *Rvi6* that originated from the Japanese crabapple *Malus floribunda* (Sieb.) sel. 821. Isolates of *V. inaequalis* able to overcome *Rvi6* have been identified in Europe, but have not yet been reported on the American continents. We recently discovered scab infection on *M. floribunda* 821 trees in a research orchard at Geneva, New York, USA, where approximately 10% of the leaves bore profusely sporulating apple scab lesions, many of which had coalesced to cover entire leaves. Chlorosis and pinpoint pitting symptoms typical of failed infections by *V. inaequalis* on hosts bearing the *Rvi6* and *Rvi7* genes were also observed. We assessed genetic diversity and population genetic structure of six *V. inaequalis* isolates collected from *M. floribunda* 821, one isolate from ‘Nova Easygro’, one isolate from ‘Golden Delicious’ and two isolates from Europe (11 isolates in total) using 16,321 genome-wide SNPs. Population genetic structure and PCA separated the isolates into distinct European and USA groups. The forgoing suggests that the new *Rvi6* virulent isolates emerged within USA populations, rather than being transported from Europe. The overcoming of resistance in *M. floribunda* 821 but not in descendant cultivars suggests that durable resistance to apple scab will require a more comprehensive understanding of *Rvi6* mediated resistance in diverse genetic backgrounds.

## Introduction

Apple scab, caused by the Ascomycete fungus *Venturia inaequalis* (Cke.) Wint., is the economically most important fungal disease of apples. In nearly all apple growing regions of the world, growers rely on fungicide applications each year to limit quality and yield loss due to scab. Scab-resistant cultivars require fewer fungicide applications, and their importance is high in organic production systems and home gardens (Brown and Maloney 2008). The commercial success of scab-resistant cultivars has been limited by less desirable fruit quality. The need to suppress scab without total reliance upon fungicides has therefore made development of scab resistant cultivars with high fruit quality one of the highest priorities of apple breeding programs.

The vast majority of the modern scab resistant cultivars originate from a Japanese crabapple clone, *Malus floribunda* (Sieb.) sel. 821, which was found to be completely free of scab in the beginning of the twentieth century. At the University of Illinois, C. S. Crandall built a collection of crabapples and, among other crossings, crossed two siblings from a *M. floribunda* 821 × ‘Rome Beauty’ progeny (Crandall, 1926). Two accessions from this cross progeny were later selected for their exceptional scab resistance yet large fruit size, and were used as resistance donors by the PRI (Purdue University, Rutgers University, and the University of Illinois) cooperative breeding program between three universities. The first PRI cultivar, ‘Prima’ was approved in 1967, and since then several breeding centers have developed resistant cultivars by crossing PRI materials (Crosby *et al.* 1990). Currently, there are more than one hundred modern scab resistant apple cultivars, with approximately 90% of them carrying the resistance derived from *M. floribunda* 821 (Brown and Maloney 2008, 2013).

To date, 18 *V. inaequalis* resistance genes (Rvi genes) including *Rvi6* and *Rvi7* from *M. floribunda* 821 and genes from other *Malus* accessions have been described. The gene-for-gene relationship has also been confirmed for the majority (Khajuria *et al.* 2018; Broggini *et al.* 2011; Bus *et al.* 2011). Based on the segregation, the resistance of *M. floribunda* 821 was also thought to rely on a single dominant gene, previously called *Vf* (syn. *Rvi6*), exhibiting resistance symptoms from hypersensitive reaction to chlorosis (occasionally necrosis or slight sporulation). However, segregation analysis of the resistance symptoms suggested the presence of an additional gene for scab resistance (Parisi *et al.* 1993; Williams and Kuc 1969). A large-scale pathogenicity study on a ‘Golden Delicious’ × *M. floribunda* 821 progeny demonstrated the presence of the *Vfh* (syn. *Rvi7*) gene responsible for the hypersensitive reaction in *M. floribunda* 821, that was distinct from the original *Rvi6* (Bénaouf and Parisi 2000). The *Vfh* gene is hypothesized to be lost early on during the breeding process, or has been overcome by the pathogen (Bénaouf and Parisi 2000; Gessler, 1989; Parisi and Lespinasse 1996).

Although the resistance granted by a single major gene can be easily inherited, as preferred by breeders, the breakdown of major genes as a result of the fast evolution of a pathogen has been documented many times in several host-pathogen systems (Pink and Hand 2003). For instance, slight sporulating symptoms were observed on ‘Prima’ (*Rvi6*) in the greenhouse trails even before its release, whereas *Rvi6* cultivars remained resistant under the field conditions. Moreover, the 1:1 segregation ratio of resistance expected from a single major gene is only observed if progeny expressing necrotic, slightly sporulating symptoms were still classified resistant (Crosby *et al.* 1990; MacHardy, 1996). In the European continent (Ahrensburg, Germany, Europe) sporulating lesions were detected on the seedlings of ‘Prima’ (*Rvi6*) for the first time in 1984, and later in 1988 on ‘Prima’, Coop 7, 9, and 10. The same inoculum was used for testing various *Rvi6* cultivars/selections, all of which were susceptible to the new isolates, but the original source of the resistance, *M. floribunda* 821, remained intact (Parisi *et al.* 1993). The breakdown of the resistance of *M. floribunda* 821 (*Rvi6, Rvi7*), was reported later in East Malling, England (Roberts and Crute 1994). Interestingly, the isolates overcoming the resistance of *M. floribunda* 821 was avirulent on ‘Golden Delicious’ and on its resistant descendants (e.g., ‘Prima’ and ‘Florina’) harboring the ephemeral *Rvi1* gene. However, it is still unclear whether the isolates were unable to overcome the resistance of these cultivars due to the *Rvi1* gene, or other genetic factors prohibited the adaptation of the pathogen (Gessler and Pertot 2012; Roberts and Crute 1994). It was reported that the background resistance of *M. floribunda* 821 is low, and its severe infection happens when the two major genes *Rvi6* and *Rvi7* have been overcome. It further suggests that the *Rvi6* resistant cultivars infected by less aggressive isolates might possess more complex resistance than that of *M. floribunda* 821 (Caffier *et al.* 2010; Parisi *et al.* 2004).

Further population genetic studies suggested that European *Rvi6* virulent isolates did not emerge from a recent mutation in the *AvrRvi6* gene, but have pre-existed for thousands of years in wild crabapple reservoirs, without relevant gene-flow to populations infecting domesticated apples. The virulent isolates on *Rvi6* cultivars are genetically less variable, but distinct from those collected from susceptible cultivars suggesting a genetic bottleneck in their recent history (Guerin *et al.* 2004; Guerin *et al.* 2007; Guerin and Le Cam 2004; Lemaire *et al.* 2015; Michalecka *et al.* 2018). Although the *Rvi6* virulent and non-virulent lineages might have been separated for thousands of years, the emergence of *Rvi6* cultivars in Europe allowed gene flow between the two lineages, which might have increased the speed of the pathogen’s adaptation, especially in orchards where susceptible and resistant cultivars coexist (Michalecka *et al.* 2018). Human facilitated transport might represent another factor for the spread of *Rvi6* virulent strains, and a possible cause of the inconsistency between the geographical origin and genetic polymorphism of isolates in several studies (Kaymak *et al.* 2016; Gladieux *et al.* 2010; Guerin *et al.* 2007; Michalecka *et al.* 2018).

New isolates that can overcome the resistance of *M. floribunda* 821 (*Rvi6* and *Rvi7* genes) have not yet been reported on the American continent. In a 33 year-long assay carried out in the Secrest Arboretum (Ohio), heavy sporulation was observed on a Japanese crabapple tree in 2003 and 2005, although its resistance stayed intact for decades. The authors however did not confirm whether the accession was identical to *M. floribunda* 821 (Beckerman *et al.* 2009). In this study, we report new *V. inaequalis* isolates collected from the infected *M. floribunda* 821 trees grown in an apple orchard, their origin and genetic diversity, and their pathogenicity on *Rvi6* resistant ‘Macfree’.

## Materials and methods

### Field assessment of apple scab symptoms

In June 2019, apple scab infection was observed on approximately 20-year old trees of *M. floribunda* 821 and scab resistant apple cultivars ‘Prima’ and ‘Nova Easygro’ grown in the research orchard Darrow Farm (Geneva, USA). In addition, this orchard has ‘Gala’, ‘Macintosh’, ‘Golden Delicious’, ‘Florina’, and four PRI cultivars. The trees were present in four replications that were distributed randomly across the entire orchard space. The orchard was maintained without any fungicide spray since 2017 (Table 2).

Apple scab incidence was assessed using the 0-9 grade scale of Lateur and Populer (1994) as follows: 0 - no observation (missing plant), 1 - no visible lesions, 2 - one or very few lesions detectable on close scrutiny of the tree (0-1%), 3 - Immediately apparent lesions in general clustered in few parts of the tree (1-5%), 4 - intermediate, 5 - numerous lesions widespread over a large part of the tree (±25%), 6 - intermediate, 7 - severe infection with half of the leaves badly infected by multiple lesions (±50%), 8 - intermediate (±75%), or 9 - tree completely affected with (nearly) all the leaves badly infected by multiple lesions (>90%). Reaction type was determined according to the scale used by the PRI breeding program (Crosby *et al.* 1990) as follows: 0 - no symptom, 1 - pin point pits, 2 - irregular chlorotic or necrotic lesions, 3 - few restricted sporulation (M) mixture of necrotic and chlorotic and sparsely sporulating lesions, or 4 - abundant sporulation.

### Sample collection

Scab infected leaves were collected from *M. floribunda* 821 and ‘Nova Easygro’ trees located in Darrow Farm (Geneva, New York) (Table 2). The fungus was isolated by sticking infected leaves to the inner top side of Petri dishes containing 1.2% water agar. The dishes were sealed and incubated overnight to let the spores fall on the agar and germinate. Germinating spores were observed under microscope and were carefully placed to new PDA plates under laminar flow to avoid the transfer of any contaminant. One scab isolate (‘VI-1797-9’) shared by H. Aldwinckle, Cornell University, Geneva, NY collected from host 4 (unspecified) in Ohio, USA (Schnabel *et al.* 1999) was also included. The well-characterized reference *V. inaequalis* isolates of European origin EU_B05 and EU_NL24 (Caffier *et al.* 2015) were obtained from INRA, France.

### Artificial inoculation of the Vf resistant commercial, cultivar ‘Macfree’

One year-old grafted plants of the *Rvi6* scab resistant ‘Macfree’ were inoculated with the *V. inaequalis* inoculum obtained from scab-infected *M. floribunda* 821 trees. Leaves were collected from each of the four *M. floribunda* 821 trees. The fresh sporulating lesions were cut, placed into a beaker containing distilled water and 0,5 μl/ml Tween 20, and shaken for 20 minutes. The suspension was filtered and the concentration was estimated by a hemocytometer, and set to at least 3×10^4^ conidia/ml by a centrifuge.

Four ‘Macfree’ scions were grafted on ‘M9’ rootstocks and maintained in a greenhouse under controlled conditions. Before inoculation, grafted plants were pruned to acquire new growth and young leaves. The spore inoculum was used to spray the first 4-6 actively growing intact leaves from the top. The inoculated plants were held in a moist chamber under 18 °C and 12h photoperiod. To maintain 100% humidity, the sprayed leaves were covered first by wet paper towels, and then by resealable plastic bags. The incubated shoot part was tagged by marking the shoots below the bags to ease future evaluation. After 48 hours of incubation, the bags were removed, and the plants were further maintained under low light, 18 °C temperature, and 70% humidity with no irrigation for 2-3 weeks (Peil *et al.* 2018). Similar to the field assessment, reaction types that were used by the PRI program (Crosby, 1990) were determined. Light microscopy was used to evaluate the intensity of sporulation on the leaves. Spores from the infected leaves were suspended in a minimal amount of water and their morphology was observed under microscope to confirm the identity of *V. inaequalis*. Images were taken using Toupview software with a digital camera for microscope (OMAX, Gyeonggi-do, South Korea) for demonstration.

### DNA extraction and genome sequencing of V. inaequalis isolates

DNA was extracted from 8 *V. inaequalis* isolates, 6 of which were obtained from *M. floribunda* 821, one from ‘Golden Delicious’, and one from ‘Nova Easygro’. In case of *M. floribunda* 821, the total 6 samples were collected from 3 different trees by sampling more than one region on two of the trees. In addition, two European isolates, EU-NL24 and EU-B05, and one U.S. isolate VI-1797-9 from Purdue were also used to extract DNA (Table 1). Cultures were grown on potato dextrose agar for four weeks. No more than 0.2g wet mycelia was cut and cleared from the agar by a sterile scalpel. Samples were ground in Eppendorf tubes with a disposable homogenizer pestle under liquid nitrogen. DNA was isolated with the Wizard Genomic DNA Purification Kit (Promega) as described by Singh and Khan (2019). The quality of DNA was checked using agarose gel (1%) electrophoresis and quantified with Nanodrop™ and UV-Vis Spectrophotometer (Thermo Fisher Scientific, USA).

**Table 1.**
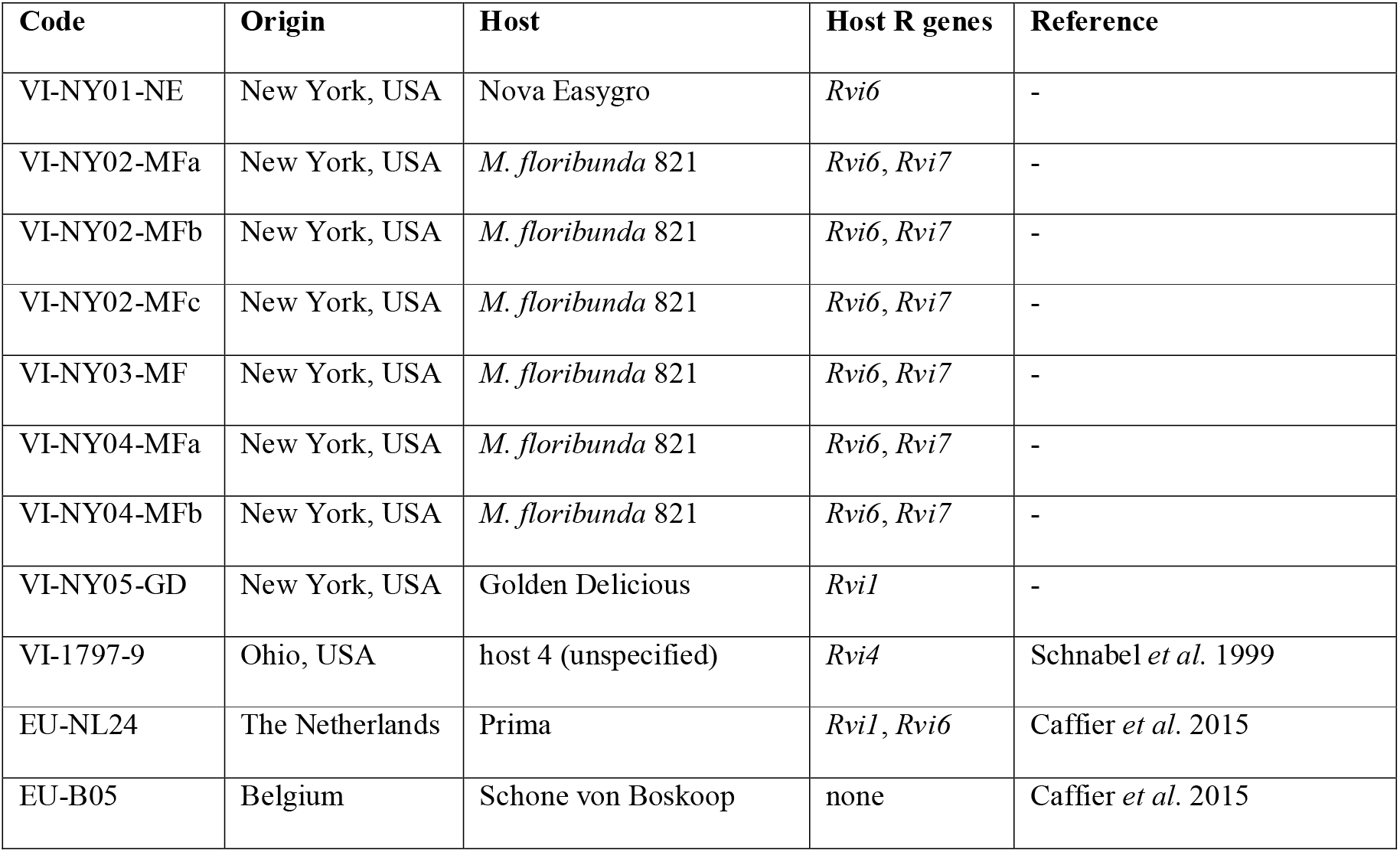
The single spore isolates of *Venturia inaequalis* originating from Europe and Darrow farm (USA) from *Malus floribunda* 821, and apple cultivars ‘Golden Delicious’ and ‘Nova Easygro’.

A total of 50ng DNA for each sample was used to prepare genome sequencing libraries for Illumina Nextera skim sequencing at Institute of Biotechnology, Cornell University, Ithaca, NY. Agilent Bioanalyzer (Agilent; www.agilent.com) was used to assess the quality and quantity of genomic libraries. Individual *V. inaequalis* samples were barcoded and sequenced in a single lane of Illumina Mi-Seq platform to generate 2×250 base pair paired-end read data.

### Sequence analysis and variant detection

Raw sequences were separated into sample-specific reads with the barcode information and used to assess the sequence quality with fastqc program (http://www.bioinformatics.babraham.ac.uk/projects/fastqc/). The program Trimmomatic (Bolger *et al.* 2014) was used to remove the sequencing adaptors and low-quality reads with quality score below 20. The genome assembly of *V. inaequalis* strain EU-B04 VeinUTG001 (Le Cam *et al.* 2019) was retrieved from National Center for Biotechnology Information (NCBI) genome database and used to align the high-quality reads from the previous step. The alignments were filtered to eliminate PCR duplicated sequences and a sorted binary alignment format (BAM) was obtained for each sample using SAMtools (Li *et al.* 2009).

Variants across the *V. inaequalis* genome were identified using Genome Analysis Toolkit (GATK version 3.8.0; McKenna *et al.* 2010) with sample-specific BAM files as input. The HaplotypeCaller plugin in GATK was used to obtain genotype variant call format (gVCF), and individual sample gVCF files were merged into a single variant file with GenotypeGVCF plugin. The raw variants were processed through “hard filtering” criteria as per GATK best practices. The resulting variants were used for base recalibration to remove false positives. The variant dataset was separated into single nucleotide polymorphisms (SNPs) and insertions/deletions (INDELs).

### Genetic diversity and population structure analysis

Genetic diversity and population genetic analysis was conducted with SNP data only. The SNPs were filtered using minimum read depth criteria of four. Furthermore, we kept SNPs with no missing data even in a single sample. Remaining high-quality SNPs were used to calculate the genome-wide statistics of diversity, population structure and selection indices in *V. inaequalis.* The population level nucleotide variation was examined through number of variants, nucleotide diversity (π), and principal component analysis (PCA) in Tassel v5 (Bradbury *et al.* 2007). The structure in *V. inaequalis* population was further examined using BYM admixture model and haploid genome option in the TESS software (Caye *et al.* 2016) with default settings. In addition, spatial coordinates were provided for the 11 isolates to infer the accurate structure in *V. inaequalis* isolates. Convergence in the population structure results was checked by running twenty independent runs for individual K values ranging from two to eight. DIC and Kmax were plotted to identify number of clusters/groups in the samples. Figures showing assignment of isolates to different clusters were generated using Clumpak - Cluster Markov Packager Across K (Kopelman *et al.* 2015).

## Results

### Apple scab infection in the orchard and under controlled conditions

In the beginning of June, scab symptoms were observed on *M. floribunda* 821 in the orchard at Darrow Farm, Geneva, NY, USA on each of the four replicated trees. The initial sparse symptoms spread to approximately 10% (incidence grade 4) of the leaves later in July and August (Table 2). The heavy infection resulted in abundant sporulation (class 4), which often covered the whole leaf surface area (Figure 1). Chlorosis and pin point pit type resistance reactions typical to the *Rvi6* and *Rvi7* genes, respectively, which are hypothetically the result of the large amount of inoculum from avirulent strains, were also frequent on the leaves.

**Figure 1.**
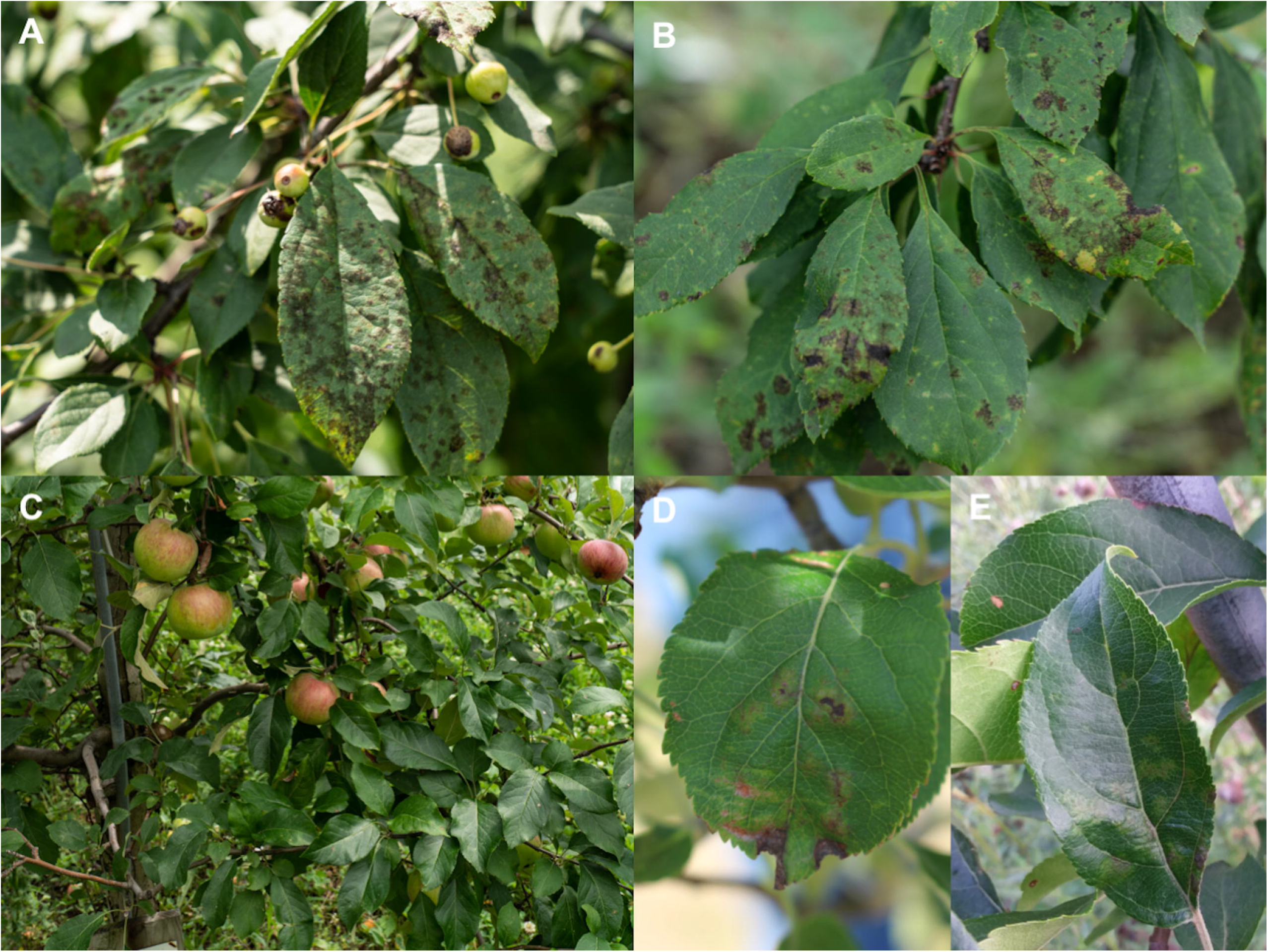
Symptoms of apple scab infection detected on *Malus floribunda* 821, ‘Nova Easygro’, and ‘Prima’: heavy sporulation (A) alongside of separate chlorotic and pin point type resistance symptoms (B) on *M. floribunda* 821; apparently intact crown of ‘Nova Easygro’ (C); weak sporulation surrounded by chlorosis and necrosis on ‘Nova Easygro’ (D) and by strong chlorosis on ‘Prima’ (E).

**Table 2.**
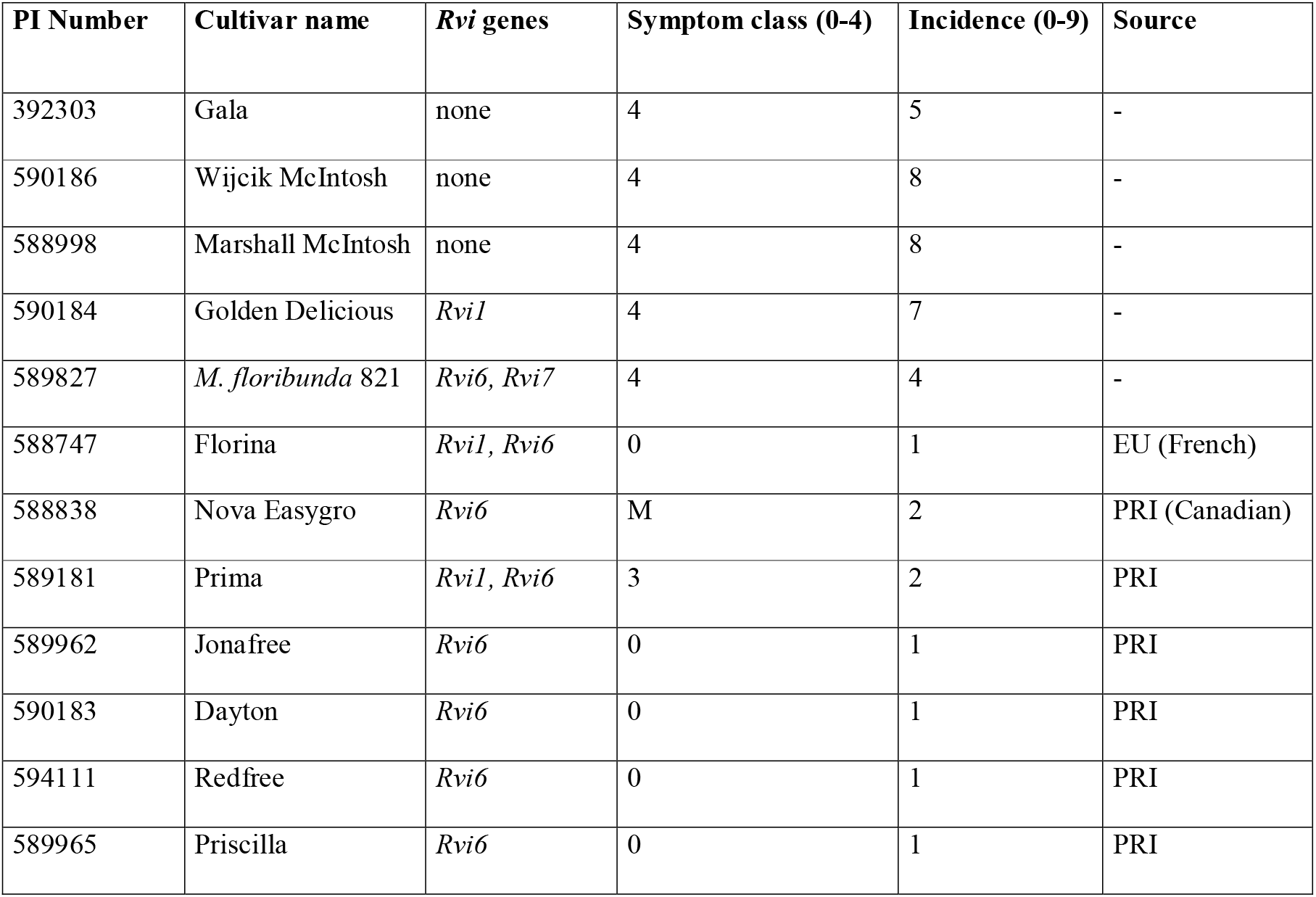
Apple scab symptoms and incidence on *Rvi6* cultivars and susceptible controls. Note: PRI stands for Purdue, Rutgers, Illinois apple breeding program.

| PI Number | Cultivar name | *Rvi* genes | Symptom class (0-4) | Incidence (0-9) | Source |
| --- | --- | --- | --- | --- | --- |
| 392303 | Gala | none | 4 | 5 | - |
| 590186 | Wijcik McIntosh | none | 4 | 8 | - |
| 588998 | Marshall McIntosh | none | 4 | 8 | - |
| 590184 | Golden Delicious | *Rvi1* | 4 | 7 | - |
| 589827 | *M. floribunda* 821 | *Rvi6*, *Rvi7* | 4 | 4 | - |
| 588747 | Florina | *Rvi1*, *Rvi6* | 0 | 1 | EU (French) |
| 588838 | Nova Easygro | *Rvi6* | M | 2 | PRI (Canadian) |
| 589181 | Prima | *Rvi1*, *Rvi6* | 3 | 2 | PRI |
| 589962 | Jonafree | *Rvi6* | 0 | 1 | PRI |
| 590183 | Dayton | *Rvi6* | 0 | 1 | PRI |
| 594111 | Redfree | *Rvi6* | 0 | 1 | PRI |
| 589965 | Priscilla | *Rvi6* | 0 | 1 | PRI |

The scab-susceptible commercial cultivars, ‘Gala’, ‘Wijcik McIntosh’, ‘Marshall McIntosh’, and ‘Golden Delicious’ all showed severe scab symptoms (Symptom class 4). Scab symptoms were more severe on ‘Golden Delicious’ than ‘Gala’ (incidence 7 and 5, respectively). Many *Rvi6* cultivars originating from *M. floribunda* 821, such as ‘Dayton’, ‘Florina’, ‘Jonafree’, ‘Priscilla’ and ‘Redfree’ were totally free of scab during the whole season. Weak symptoms were also detected on the descendants of *M. floribunda* 821, ‘Prima’, and ‘Nova Easygro’; however, on these cultivars the infection was barely detectable. No more than one or two leaves per tree were infected in these cultivars (grade 1). In case of ‘Nova Easygro’, both chlorosis and necrosis were detected around the slight sporulation as the sign of induced defense response (M), while the weak sporulating lesions on ‘Prima’ were all surrounded by unusually strong chlorosis (3).

Abundant sporulating scab symptoms were observed under microscope on the leaves of ‘Macfree’ after 15 days of inoculation in controlled environment (Figure 2). The sporulation covered a large part of the leaf surface area without the trace of restriction by chlorotic or necrotic response, hence we consider this reaction to represent the full compatibility infection class (4).

**Figure 2.**
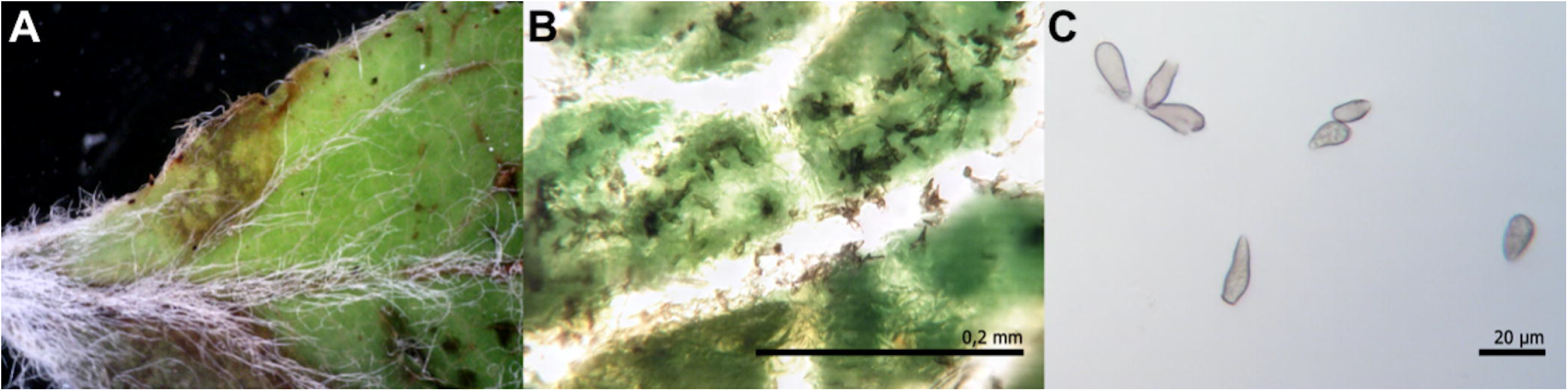
Susceptible apple scab reaction of ‘Macfree’ (Rvi6) in a controlled environment to the inoculum of *Venturia inaequalis* isolates collected from the field from *Malus floribunda* 821. Sporulating leaf surface at 16× magnification (A), macroscopic lesions (B), and suspended conidiospores at 40× magnification (C).

### Genomic diversity in the Venturia inaequalis isolates

To understand the genetic diversity, population structure and admixture, we sequenced and analyzed the 11 *V. inaequalis* genomes in this study (Table 1). A total of 34.5 million raw sequencing reads were obtained, out of which 0.38% low quality reads were discarded for further analysis (Table 3). The filtered sequence data constitutes about 3.1 million average reads per sample, and provided 9.2X coverage of the *V. inaequalis* genome in NCBI. The read alignment rate against the *V. inaequalis* genome varied from 97.7% to 99.2%, with an average of 98.7% (Table 3).

**Table 3.** Summary statistics of number of sequence reads generated, quality filtering, and percent alignment (mapping) against the *Venturia inaequalis* reference genome (Le Cam *et al.*, 2019).

| Sample ID | Raw Reads | Cleaned Reads | Mapping (%) |
| --- | --- | --- | --- |
| VI-NY05-GD | 3549542 | 3549542 | 99.08 |
| VI-NY01-NE | 3223354 | 3223354 | 98.91 |
| VI-NY03-MF | 2850184 | 2850184 | 99.1 |
| VI-NY04-MFa | 3342636 | 3342636 | 99.2 |
| VI-NY04-MFb | 2453088 | 2453088 | 98.74 |
| VI-NY02-MFa | 3128934 | 3128934 | 98.93 |
| VI-NY02-MFb | 2770586 | 2770586 | 97.69 |
| VI-NY02-MFc | 2602972 | 2602972 | 99.27 |
| VI-1797-9 | 4555028 | 4509802 | 98.75 |
| EU-NL24 | 3015184 | 2973782 | 98.41 |
| EU-B05 | 2990158 | 2945712 | 98.54 |
| **Total** | 34481666 | 34350592 | 98.7 |
| **Coverage** | 9.21 |  |  |

After variant detection and GATK hard filtration, total of 199,607 SNPs were identified with no missing data and representing one SNP every 360 base pairs of the *V. inaequalis* genome. About 38.1% (n=76,130) SNPs were detected in European isolates and 25.2% (n=50,292) were present in the two isolates collected from U.S. locations other than Darrow farm (Supporting Figure S1). The seven isolates from Darrow farm constitute about 68.3% (n=136,406) of total SNPs identified. Approximately 4.4% of SNPs were shared between these three groups of *V. inaequalis* isolates (Supporting Figure S1). The 6 isolates from *M. floribunda* 8221 had 119,109 SNPs representing about 59.6% of total genetic diversity in the population. There were 55,130 SNPs identified in the three replicated isolates collected from the same *M. floribunda 821* tree. We further observed about 0.29 nucleotide diversity (π) in this *V. inaequalis* collection. The seven isolates collected from Darrow farm had a π value of 0.39 and 6 isolates from *M. floribunda 821* exhibited π value of 0.43, thus representing slightly higher nucleotide diversity than the entire collection. The nucleotide diversity of the two U.S. isolates outside of Darrow farm and the two European isolates was not estimated due to limited sample size.

### Population Genetic Structure Analysis

The population structure was examined using 16,321 high-quality SNPs to perform principal component analysis (PCA) and BYM admixture estimation of *V. inaequalis* genomes. PCA clearly separated the 11 isolates into two distinct European and U.S. groups (Figure 3). The two European isolates from Belgium and Netherlands also showed clear distinction from each other. The U.S. groups also exhibited slightly dispersed pattern (Figure 3). For instance, two *V. inaequalis* isolates from outside the Darrow farm had moderate separation from the remaining isolates. Five of the seven isolates collected from Darrow farm displayed a close clustering, while two of them had a distinct placement on PCA biplot. The latter two isolates represent two of the three replications that were collected from the same tree of *M. floribunda* 821. The remaining isolate co-localize with the other isolates from *M. floribunda* 821. The other subgroup of 5 U.S. isolates from Darrow farm had diverse host range including ‘Golden Delicious’, ‘Nova Easygro’, and *M. floribunda* 821.

**Figure 3.**
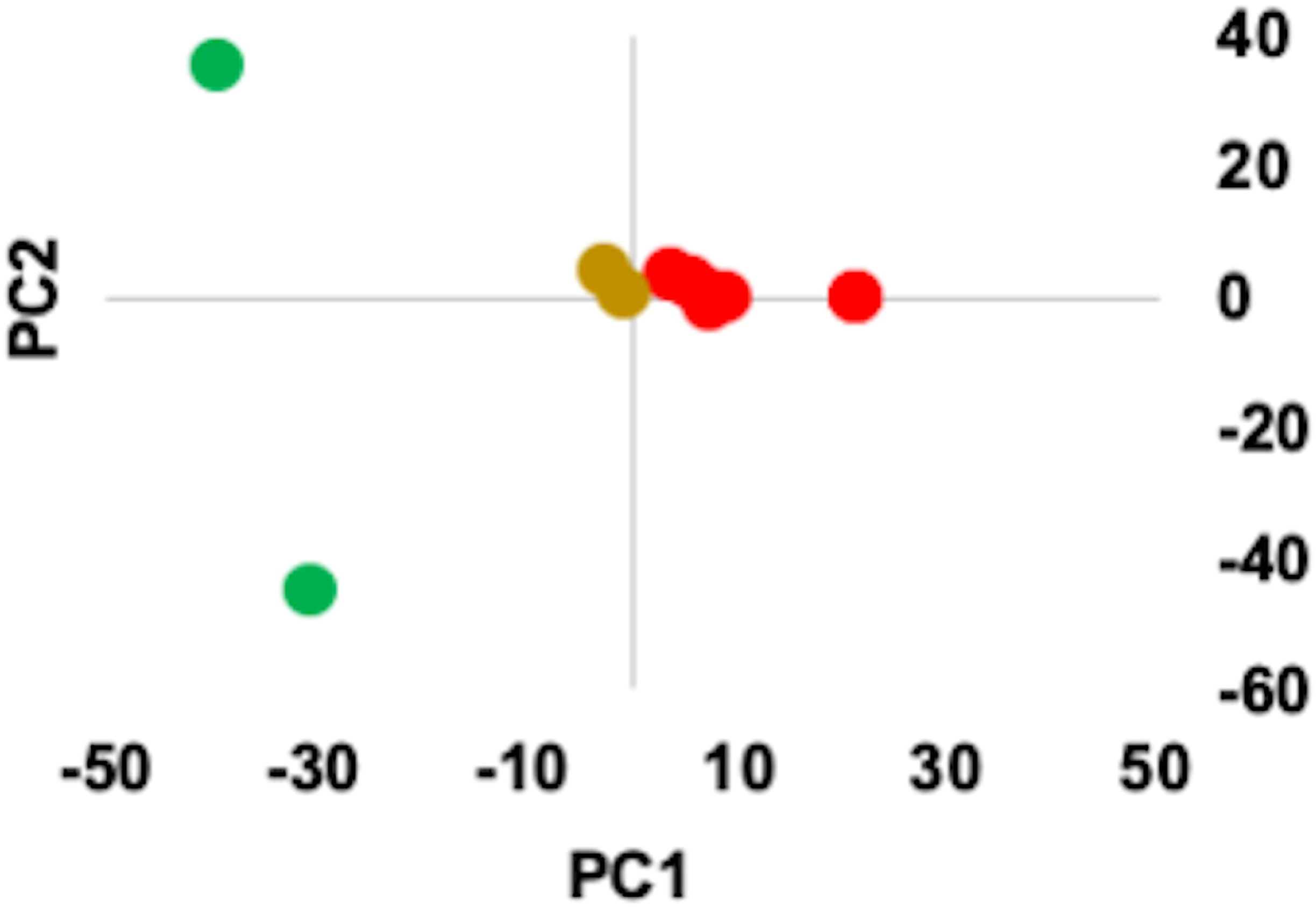
Principal Component Analysis of eleven *Venturia inaequalis* isolates using 16,321 SNPs supported by a minimum of four sequencing reads. The original variant dataset was filtered to retain SNPs, supported by minimum four sequencing reads. The color of points on PC A biplot represent different isolates: European (green), American from *Malus floribunda* 821 (red), American from ‘Golden Delicious’ and ‘Nova Easygro’ (yellow).

A genome admixture analysis reflected four main clusters across the 11 *V. inaequalis* isolates (Figure 4). Two of the clusters represent the two European isolates and the other two clusters were specific to the U.S. isolates. A major proportion of genomic composition was different between the two European isolates, which also differed from the remaining isolates from U.S. regions. A U.S. isolate VI-NY05-GD obtained from ‘Golden Delicious’ exhibited partially different genomic composition than the other U.S. isolates from within and outside Darrow farm. The six isolates from *M. floribunda* 821 mostly showed similar genome composition except VI-NY03-MF, which had some admixture from European genomes (Figure 4). The main differences observed in genomic composition at K=4 remained consistent as K value changed from 3 to 5. At K=5, the few U.S. isolates and two European isolates showed a small proportion of distinct genome other than the four groups observed at K = 4.

**Figure 4.**
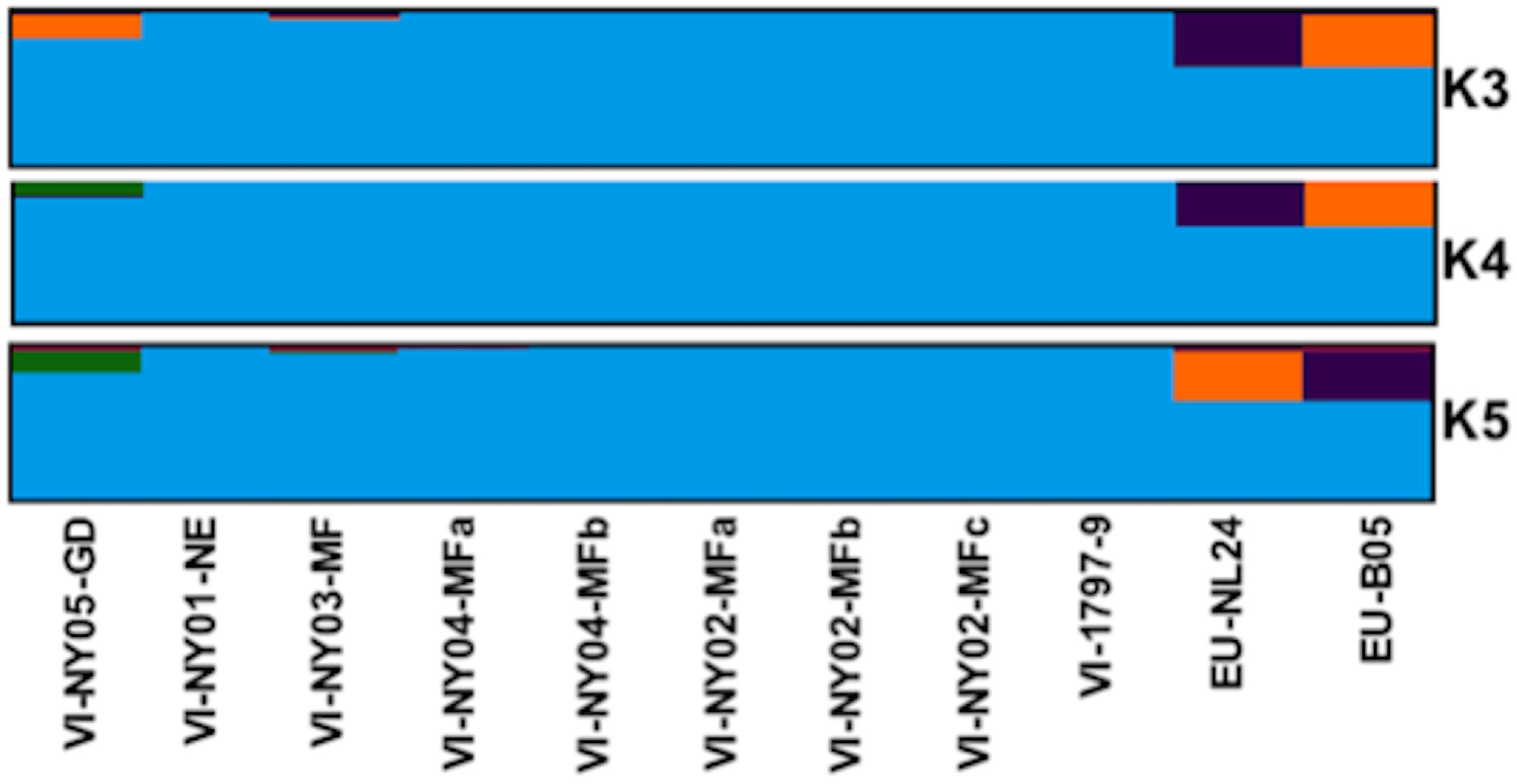
Plot showing population structure and admixture levels in the genomes of 11 *Venturia inaequalis* isolates using 16,321 SNPs supported by minimum four sequencing reads. The analysis was conducted with the TESS software (Caye *et al.* 2016) using BYM admixture model and haploid genome with default settings. Convergence in the population structure results was checked by running twenty independent runs for individual K values. The best population structure was represented by K=4.

## Discussion

We observed severe apple scab symptoms on trees of the Japanese crabapple *M. floribunda* 821, which carries the scab resistance genes *Rvi6* and *Rvi7*, in a research orchard at Geneva, New York, USA in 2019. The breakdown of *Rvi6* resistance was reported within a decade of its deployment from several European locations (Bénaouf and Parisi 1997; Parisi *et al.* 1993), but until now resistance has remained effective elsewhere. This is the first confirmed loss of the resistance of *M. floribunda* 821 in North America. Seven cultivars descended from *M. floribunda* 821 (‘Dayton’, ‘Jonafree’, ‘Prima’, ‘Priscilla’, and ‘Redfree’, ‘Florina’, and ‘Nova Easygro’) within the same orchard developed levels of scab much lower than observed on Japanese crabapple, despite their common ancestry with Japanese crabapple and possession of *Rvi6* (Table 2). The high incidence of scab across a range of known susceptible cultivars, and the uniform infection among all replicate trees of *M. floribunda* 821, would indicate that low scab incidence among these remaining resistant cultivars was not due to escape, but because of the presence of *Rvi6* in a genetic background still contributing additional resistance to *V. inaequalis.* The foregoing would be consistent with the report by Roberts and Crute (1994) that many *Rvi6* cultivars remained resistant to scab in Europe after the resistance of *M. floribunda* 821 had eroded there. Parisi *et al.* (2004) also reported that ‘Prima’ remained more resistant to scab than *M. floribunda* 821 when challenged by isolates able to overcome *Rvi6.* It has been proposed that *Rvi1*, derived from ‘Golden Delicious’, might contribute additional resistance when paired with *Rvi6*. However, we observed continued resistance to scab among many *Rvi6* cultivars that are not known to carry *Rvi1*, meanwhile slight sporulation was observed on ‘Prima’ carrying *Rvi1* alongside *Rvi6* (Table 2). Gessler and Pertot (2012) speculated that *Rvi1* might occasionally act indirectly against *Rvi6* virulent isolates, and Caffier *et al.* (2010) hypothesized a fitness cost to virulence towards *Rvi6* conferred by loss of *avrRvi6.* However, the ability to overcome a larger number of R genes (excluding *Rvi6)* has not correlated with low fitness in past studies (Parisi *et al.* 2004; Peil *et al.* 2018).

It is important to distinguish between mild chlorotic flecking typical of greenhouse inoculations and the loss of field resistance to apple scab, wherein profusely sporulating lesions are macroscopically visible in orchard assessments. The chlorotic and necrotic symptoms seen on the field on ‘Prima’ and ‘Nova Easygro’ cannot be considered as evidence that their field resistance has been overcome. Likewise, chlorotic lesions supporting sparse sporulation have been reported on ‘Prima’ following greenhouse inoculations (Crosby *et al.* 1990). Even in cases of compatible reaction between the host and the pathogen, scientific agreement on whether resistance can be considered broken might depend on the severity and economic impact (Delmotte *et al.* 2016). Sparse chlorotic sporulation on commercial *Rvi6* cultivars is not generally considered a sign of resistance breakdown (Bus *et al.* 2011). On the other hand, *Rvi6* cultivars including ‘Macfree’ often appear to be susceptible under greenhouse conditions, indicating the breakdown of *Rvi6* resistance, but remains highly resistant in field plantings (Crosby *et al.* 1990; Roberts and Crute 1994). We observed similar mild and chlorotic symptoms on several scab resistant cultivars in the field (Table 2), but these are easily distinguished both qualitatively and quantitatively from the symptoms recorded on *M. floribunda* 821.

The earlier discovery of *Rvi6* and Rvi7-virulent isolates in Europe creates the possibility of transportation of such isolates to North America and subsequent spread. However, our comparison of genetic polymorphism of the European and American virulent isolates suggested that both evolved or were selected from genetically distinct and separate populations. In Europe, the *Rvi6* virulent lineage has existed separately in non-agricultural areas for thousands of years (Lemaire *et al.* 2015). While there is considerable genetic distinction between the isolates from ‘Golden Delicious’ and the new virulent isolates, we do not have sufficient data to entirely exclude the possibility of gene flow and human transportation of isolates.

The considerable genetic polymorphism among the *M. floribunda* 821 virulent American isolates makes them less likely to be clonal variants. Interbreeding between *Rvi6* virulent and non-virulent isolates has already been documented in mixed orchards (Michalecka *et al.* 2018). Even though our isolates were collected within an unsprayed experimental orchard comprised of several cultivars, genetic dissimilarity detected between the groups suggests that intense hybridization has not yet occurred between the *M. floribunda* 821 virulent and avirulent isolates studied here.

To successfully manage the disease resistance of cultivars, the distribution and evolutionary processes of pathogen isolates must be understood. The breakdown of resistance from *M. floribunda* 821, but not in descendant cultivars, suggests that the overall genetic background into which a single major R gene is incorporated can substantially affect outcomes with respect to durability of resistance over time. Broad-spectrum and partial polygenic resistance is considered a durable alternative to major resistance genes (Corwin and Kliebenstein 2017; Robinson, 1996; Khan and Chao 2017). Breeders prefer to choose the most promising disease resistance genes and gene combinations to achieve commercially successful cultivars with durable disease resistance. However, due to the time needed to select and breed for polygenic resistance and the possibility of linkage drag of undesirable alleles, breeding programs have prioritized major monogenic resistances. In-depth understanding of the factors resulting in the emergence of *M. floribunda* 821 resistance-breaking isolates, and the breakdown of resistance from *M. floribunda* 821 but not in descendant cultivars, may ultimately reduce the risk of eroding the resistance of *Rvi6* cultivars and enhance their durability.

## Acknowledgments

This research was supported by the New York Apple Research and Development Program (ARDP). We thank David Strickland and Liqiang Gao for culturing and DNA extraction of some of the isolates used in the study and Della Cobb-Smith for technical support.

---

**Supporting Figure S1**. Venn diagram showing the distribution of total SNPs in *Venturia inaequalis* isolates from Europe and USA (both Darrow farm and non-Darrow farm).

