## Supplementary figures and images for "New North American isolates of *Venturia inaequalis* can overcome apple scab resistance of *Malus floribunda* 821"

### Supporting Figure S1

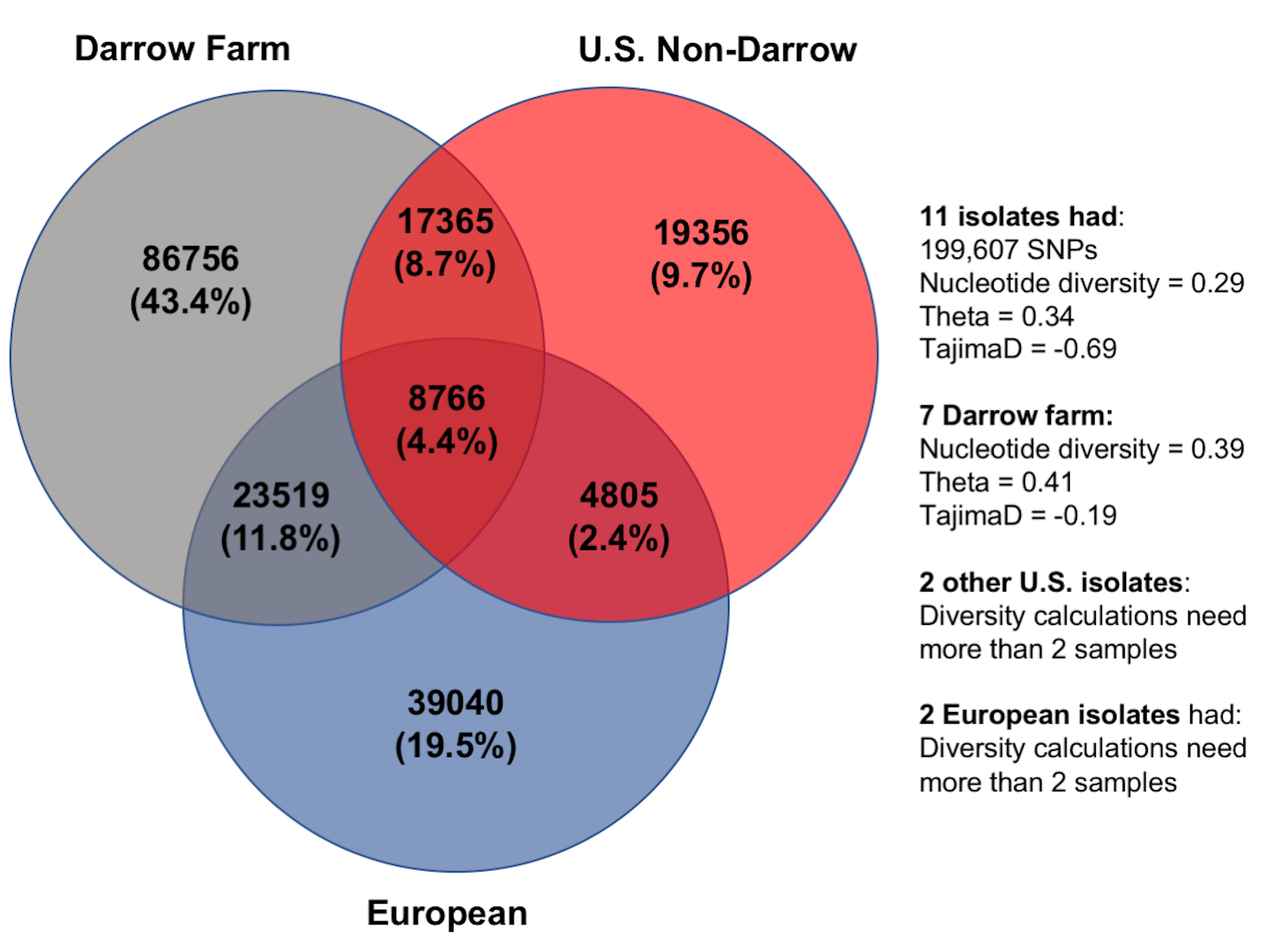
